# A Triple-Hit Alcoholic Liver Disease Model Linking High-Fat Diet, Endotoxins, and Neo-Antigen Formation

**DOI:** 10.1101/2025.11.10.687007

**Authors:** Shivani Arora, Anju Bansal, Anju Katyal

## Abstract

**BACKGROUND:** Existing rodent models of alcoholic liver disease (ALD) often fail to replicate its full clinical progression—from steatosis to alcoholic hepatitis and ultimately cirrhosis—and overlook key metabolic contributors. Clinically, high-fat diets and gut-derived endotoxins (from enteric dysbiosis) act as additional “hits” that exacerbate alcohol-induced liver injury. Moreover, alcohol metabolism generates oxidative stress and modifies self-proteins into neo-antigens, potentially triggering immune responses. A model that incorporates these elements is critical for studying ALD pathogenesis and its immunological aspects.

**METHODS:** Male Wistar rats were randomized into four groups. The AL group received alcohol (36% caloric equivalent) and a single LPS dose (1 µg/kg) 24 hours before sacrifice. The AF group was fed a high-fat diet (35% caloric equivalent) for 15 days, then maintained on alcohol and fat. The ALF group was similarly primed with fat, maintained on alcohol and fat, and given LPS prior to sacrifice. The ICC group served as isocaloric controls. Liver injury was assessed via histology, biochemical markers (AST, ALT, MDA), and serum endotoxin levels. Antibody titers against liver proteins were analyzed by ELISA; neo-antigens were identified using proteomics.

**RESULTS:** All treated groups showed elevated liver enzymes, MDA, and endotoxin levels, indicating progressive liver damage. The ALF group exhibited the most severe pathology within six weeks, progressing from steatosis to steatohepatitis and fibrosis. In contrast, the AF group showed steatosis alone, and the AL group displayed moderate inflammation. Neo-antigens were most abundant in the ALF group.

**CONCLUSION:** We present a novel, rapid-onset triple-hit model of ALD that mimics the disease’s full spectrum. This model enables mechanistic studies on how diet and gut-derived endotoxins contribute to ALD and highlights the immunogenic potential of alcohol-induced neo-antigens.

## INTRODUCTION

Alcoholic liver disease (ALD) is recognized as the major health threat worldwide. Epidemiological statistics reveal that only 10 to 15% of chronic alcohol abusers ever develop cirrhosis ^1^. This presses strongly on the fact that alcohol alone is not solely responsible for initiation and/or progression of ALD and demonstrates the significance of predisposing or disease promoting factors which aggravate the condition. In the development of ALD; interaction of primary (alcohol and its metabolites) and secondary risk factors (dietary habits, endotoxins, co-morbid diseases, genetic predisposition and gender) hasten the disease succession^2, 3^. Of many secondary risk factors-high fat diet, and gut derived endotoxins are the ones known to play a definative role in development of steatosis and propagation of inflammation during development of ALD ^4, 5^. Chronic obesity has been evidenced to sensitize liver by activating innate immune responses initializing fatty liver syndrome, and chronic alcohol consumption has been evidenced to damage gut barrier integrity, pouring gut derived endotoxins in serum that acts as a direct priming signal for macrophage polarization.

Where complex patho-mechanisms of ALD have been well investigated, we still need an animal model that recapitulates the complexity of disease within a reasonable time frame. However, these models of ALD suffer with a major drawback of being unable to produce end stage liver disease. The most commonly used animal model for alcoholic liver disease employs alcohol administration (upto 36% of caloric equivalent) in Lieber de Carlie diet, is although easy to perform, results either in alcoholic steatosis or requires exceptionally long period of chronic alcohol exposure of more than 20 weeks in rodents fibrosis^15, 16^. This model also fails to achieve consistently high blood alcohol levels which is a prerequisite for alcohol induced liver damage^17^. To counter this failure of achieving end stage liver pathology ‘**Two/Double Hit** strategy that combines the use of a cofactor along with alcohol is frequently adapted. One such widely accepted model is an intragastric alcohol infusion model developed by Tsukamoto et al. this model employs co-administration of high fat diet and alcohol for 120 days to produce high blood alcohol levels, focal necrosis and mild fibrosis, though this model produces progressive liver injury, is both time consuming as well as requires surgical expertise, and even has high mortality rate^11, 12^. Further, in this model the role of LPS, which has been clearly implicated in pathogenesis of alcoholic cirrhosis and chronic active hepatitis ^18, 19^ is overlooked. To overcome this Koteish et al employed LPS in their studies along with alcohol to set-up long term rodent models for analyzing the interactions between alcohol & LPS ^6, 7, 8^. Another popular and widely used animal model-the NIAAA/Gao-binge model-uses a mix of binge and chronic ethanol administration for 10 days-has proven to be extremely useful in understanding the diversity and interplay between different immune cell populations. These existing animal models have contributed significantly to understanding the input of secondary risk factors individually in development of ALD, but still fail to achieve the typical end stage disease patterns for ALD unless overtly long periods of study are employed. Therefore, to have a clear picture on how fat and gut derived endotoxins; together affect the progression of alcoholic liver disease it is imperative to create a “Triple Hit” model for alcoholic liver disease. Taking lead from our preliminary data for the current study we hypothesized that by using a combination of 3 hits-High Fat Diet (First Hit) as sensitizing agent; Alcohol as the Second/Direct hit; and Endotoxins (administered as single shot of LPS at the end of the study) as the Third Hit to polarize immune cells; we can create a progressive model for alcoholic liver disease that will display the entire spectrum of disease starting from fatty liver and culminating into liver cirrhosis within a short period 10 weeks. For this purpose, we maintained Male Wistar rats on either alcohol plus LPS, or on alcohol plus fat or on alcohol plus fat plus LPS for duration of six (42 days), eight (56 days), and ten weeks (70 days). Simultaneously three groups of animals were maintained on an isocaloric diet for equivalent duration which served as control. In all the groups, we performed histological analysis and liver function tests to track the progression of liver damage. The Blood alcohol levels (BALs), serum endotoxin levels and Malondialdehyde (MDA) levels were also monitored as parameters of alcohol intoxication, hepatic injury and lipid peroxidation respectively.

Further, in this study we monitored humoral immune responses in terms of serum antibody titter against liver proteins using ELISA and established the identity of neo-antigens formed during the progression of disease using peptide mass fingerprinting.

## MATERIAL and METHODS

### Materials

Ethanol (Emsure, Merck Chemicals), LPS (E. Coli L8274: S. Enterica L6011 used in ratio of 1:1, Sigma, USA), Corn oil (Cornola, Mawana Sugars, Haryana, India), Isocaloric control diet (Rodent Liquid Diet-LD 102, TestDiet, St. Louis, MO, USA), Test diet with alcohol (Alcohol Rodent Liquid Diet-LD 102A, TestDiet, St. Louis, MO, USA)

### ANIMALS

Male Wistar rats (6 to 8 weeks old, 120 to 130g) were procured from the Dr. B.R. Ambedkar Centre for Biomedical Research animal house facility and all the procedures were approved by the ethical committee. The animals were maintained at 27±2°C for a 12-h dark and light cycle. Animals had free access to the feed and water. Each animal was maintained in a separate cage and were acclimatized for a period of 7 days before starting the treatment. Anthalmintic treatment (ivermectin 0.4mg/kg) was given once weekly for 2 weeks prior to the start of experiment. All the experimental procedures were performed according to guidelines for Care and Use of Laboratory Animals approved by CPCSEA and Institutional Animal Ethics Committee.

### Experimental Design

There were 12 groups which received test and control diets as per the experimental schedule; the type of treatment in respective doses is given in table# 1. The animals were fed with these combinations for 6, 8 & 10 weeks. In alcohol and LPS group the animals were administered ethanol (accounting for 36% of total calories), and received a single dose of LPS I.V. 24 hrs prior to the sacrifice. In the alcohol and fat group, the animals were maintained on fat (equivalent to 35% of total calories) for first 15 days and were then administered alcohol (equivalent to 36% of total calories) and fat (35% caloric equivalent). In the case of alcohol, fat and LPS group, the animals were maintained on test diet containing fat (35% of caloric equivalent) for first 15 days and their after were maintained on ethanol (36% of caloric equivalence) and fat (35% of total calories). The animals were anesthetized with pentobarbital-sodium 50 mg/kg and blood was collected to obtain serum. Following this, livers were harvested and perfused with 0.1M PBS (pH7.2). The harvested livers were stored in liquid nitrogen. A portion of liver was preserved in a solution of 10% formal saline to perform histopathology.

**Table No. 1:** Treatment Schedule for different groups.

| Group | Treatment | No. of Animals | Duration (Weeks) |
| --- | --- | --- | --- |
| 1 | Isocaloric control diet (ICC group) | 8 | 6 |
| 2 | Isocaloric control diet (ICC group) | 8 | 8 |
| 3 | Isocaloric control diet (ICC group) | 8 | 10 |
| 4 | Alcohol + LPS (AL group) | 8 | 6 |
| 5 | Alcohol + LPS (AL group) | 8 | 8 |
| 6 | Alcohol + LPS (AL group) | 8 | 10 |
| 7 | 15 Days Fat followed by Fat + Alcohol (AF group) | 8 | 6 |
| 8 | 15 Days Fat followed by Fat + Alcohol (AF group) | 8 | 8 |
| 9 | 15 Days Fat followed by Fat + Alcohol (AF group) | 8 | 10 |
| 10 | 15 Days Fat followed by Fat + Alcohol + LPS (AF group) | 8 | 6 |
| 11 | 15 Days Fat followed by Fat + Alcohol + LPS (ALF group) | 8 | 8 |
| 12 | 15 Days Fat followed by Fat + Alcohol + LPS (ALF group) | 8 | 10 |

### HISTOPATHOLOGY

Livers fixed in fixed in 10% formalin were embedded in paraffin for preparation of histological sections. 5µm thick sections were cut, and hematoxylin and Eosin, Masson trichrome as well as reticulin staininbg was performed to analyze inflammatory changes and fibrosis. Histopathology was done in a blind manner by an independent pathologist.

### Alanine transaminase (ALT) & Aspartate transaminase (AST) Levels in Liver Tissue

AST and ALT levels were measured in the serum, for these kits from Autopark Siemens were used. The kits used aspartate and alanine as substrate, the rate of consumption of NADH to NAD^+^ was measured kinetically, by recording the decrease in absorbance at 340nm.

### Blood Alcohol Content Profile

A single bolus dose of 40% v/v (2g/kg body weight) ethanol was given to rats of all the groups before starting the experiment. To determine alcohol metabolism, blood was withdrawn from the tail vein at an interval of 0 min, 30 min, 60 min, 90 min and 120 min in heparanized tubes, centrifuged at 2000g to collect plasma. Ethanol concentration in plasma were estimated by measuring absorbance at 565nm resulting from coupling of NADH with formazan MTT using EnzyChrom kit (ECET 100, Bioassay Systems, Hayward, CA, USA).

### Endotoxin Levels

A colorimetric based method was used for detection of lipopolysaccharides in the sample (GenScript USA), in which the endotoxin catalyzes the activation of proenzyme in limulus amoebocyte lysate, that ultimately cleaves a colorless substrate to produce a colored end-product. This end product was measured spectrophotometrically at 545nm. The concentration of endotoxin in the test sample is extrapolated from the standard curve made using different dilutions of lyophilized endotoxin standard (1EU/ml ≈ 0.1ng/ml).

### Liver Tissue MDA Levels

Malondialdehyde formation in each sample is taken as a measure of lipid peroxidation and is a marker for oxidative stress. It was measured spectrophotometrically following the protocol of Ohkawa et al, 1979^13^. Liver tissue homogenate (equivalent to 100µg of protein) was mixed with 1ml of 20% acetic acid (pH 3.5), 1ml of 0.67% thiobarbituric acid and 0.1ml of 8% SDS. The mixture was kept in a water bath at 100 °C for one hour. The pink chromogen formed in the reaction mixture was extracted with n-butanol and the absorption was measured at 535nm. The results were expressed as nmol/hr/mg protein of MDA formed using a molar extinction coefficient of 1.56X 10^5^ M^-1^ cm^-1^.

### 4.10. Liver Homogenization and Subcellular Fractionation

Livers were homogenized in 10 volumes of ice cold homogenization buffer (10mM Tris-HCl, pH-7.5, 250mM sucrose, and protease inhibitor cocktail (Sigma Aldrich, USA) to obtain the crude protein preparation. The protein concentration was estimated using Bradford’s Assay (Bradford, 1976). The liver homogenates were centrifuged at 1000g for 10 minutes at 4°C to remove cell debris and nuclear fraction from pellets. The supernatants were again centrifuged at 10,000g for 20 minutes at 4°C to obtain mitochondrial fraction in the pellet and the supernatants from step 2 were subjected to ultracentrifugation at 100,000g for 1 hour at 4°C to pellet down Microsomal fraction. The supernatant thus obtained was used as cytosolic protein fraction. The protein content was quantitated using Bradford assay.

### Investigation of Antibody Response By ELISA

96 well polystyrene microplates for ELISA (Maxisorp, Nunc) were coated overnight with 2µg of protein from the cytosolic, microsomal and mitochondrial fractions at 4°C in carbonate bicarbonate buffer (0.05M, Ph 9.6). After incubation free unbound antigen was washed with TTBS (TBS-0.05%Tween 20), and then blocked with 1%BSA in TTBS for 1hour at 37°C. The plates were again washed with TTBS. The serum from control, and experimental rats was diluted to 1:100 with blocking buffer and added in triplicate to respective wells and incubated for 2 hours, followed by washing with TTBS and incubation with secondary, goat polyclonal anti-rat IgG HRP tagged antibody (1:5000, ab 102187, Abcam), goat polyclonal anti-rat Ig A HRP tagged antibody (1:10,000, ab 97185, Abcam) and mouse monoclonal anti-rat Ig E Biotin tagged antibody (1:5000, ab 11666, Abcam) for 2 hours. For detection of IgE response, further incubation with streptavidin HRP (1:10,000, sc 52234, Santa Cruz Biotechnology, Inc.) was performed for 1 hour. Finally after washing with TTBS, the reaction was developed using OPD (ortho-phenylene diamine) for 15 min and reaction was stopped using 4N H_2_SO_4_.

### 4.16. Two-Dimensional Gel Electrophoresis and Immunoblotting

The cytosolic protein fractions of livers from control and alcohol treated animals (Group 1-12) were precipitated using 2D protein precipitation kit from Geno Biosciences. The protein pellet was dissolved in rehydration buffer (50mM DTT, 1% ampholytes, protease inhibitor cocktail, PMSF and tracking dye). Each experiment was performed in duplicate. 2mg protein was loaded onto 11cm Immobilized pH gradient (IPG) strip, pH 3-10 nl (non linear) from Serva (Heidelberg, Germany). The IPG strips were rehydrated at room temperature for 16 hours, equilibrated in equilibration buffer I (50mM DTT) and II (40mM Iodoacetamide) for 30 minutes each and focused using Iso Electric Focusing Cell (Protean IEF Cell, Biorad, CA), by a standard program initiated with 250 V for 15 min, followed by linear ramping at 1000 V for 2 Hours and final focusing at 8000V for a total of 40,000 VH. The second dimension of electrophoresis was performed on 12% SDS polyacrylamide gel. Each protein sample was processed in duplicate. One of the gels was used for immunoblotting; the proteins from the gel were transferred onto the nitrocellulose membrane (0.45µm, Advanced Microdevices Pvt. Ltd., India) using semidry transfer blot (Bio Rad). The membranes were blocked with 3% BSA in PBS for 1hr, after 3 washings with PBST buffer, the membranes were probed overnight with the primary antibodies (sera of rats) (dilution 1: 50 in PBST) at 4°C. The blots were then washed thrice with PBST and incubated in secondary antibody; sheep polyclonal anti-rabbit IgG HRP tagged antibody (1:5000in PBST, Abcam); goat polyclonal anti-rat IgG HRP tagged (1:2500 in PBST, Abcam); goat polyclonal anti-rat IgA HRP tagged (1:5000 in PBST, Abcam); mouse monoclonal anti-rat IgE biotin labeled (1:2500 in PBST, Abcam) for 2 hours at room temperature (for detection of IgE reactive proteins one more step of incubation with streptavidin-HRP was incorporated before developing the membrane using chemiluminescence). After 3 washings with PBST the reactive proteins were identified using chemiluminescence kit (Amersham). The second gel was stained with comassie brilliant blue stain to serve as a 2D reference map for protein spot identification. The spots were indexed using PDQUEST software from Bio Rad.

### 4.18. Protein characterization

CBB stained protein bands were excised from the corresponding SDS-polyacrylamide gels. The gel slice was diced to small pieces and placed in sterile eppendorf tubes. The gel pieces were destained using destaining solution for 10minute intervals (3-4 times) until the gel pieces become translucent white. The gels were dehydrated using acetonitrile and Speedvac till complete dryness. The gel pieces were rehydrated with DTT and incubated for one hour. After incubation the DTT solution was removed. The gel pieces were now incubated with Iodoacetamide for 45min. The supernatant was removed and the gel was incubated with ammonium bicarbonate (100mM) solution for 10min. The supernatant was removed and the gel was dehydrated with acetonitrile for 10min and Speedvac till complete dryness. Trypsin (T6567, Sigma) solution was added and incubated overnight at 37^0^C. The digested solution was transferred to fresh eppendorf tube. The gel pieces were extracted thrice with extraction buffer and the supernatant was collected each time into the eppendorff above and then Speedvac till complete dryness. The dried peptide mix was suspended in TA buffer. The peptides obtained were mixed with HCCA matrix in 1:1 ratio and the resulting 2ul was spotted onto the MALDI plate. After air drying the sample, it was analyzed on the MALDI TOF/TOF Brukers Daltonics ULTRAFLEX III instrument and further analysis was done with FLEX ANALYSIS SOFTWARE for obtaining the PEPTIDE MASS FINGERPRINT. The masses obtained in the peptide mass fingerprint were submitted for Mascot search in NCBI nr database for identification of the proteins.

### Statistics

All the experiments were repeated 3 times. Data was analyzed using Origin 7.0 software and graphs were made in Microsoft office Excel 2007. All the results were expressed as Mean ± S.E.M. and analyzed using one way ANOVA followed by student t-test; p < 0.001, p < 0.01 and p < 0.05 were considered statistically significant.

## Results

**H**&E staining and Masson trichrome staining were performed to track morphological changes in liver after chronic alcohol administration. In the treated animals, progressive increase in hepatic injury was observed which was dependent on the type and duration of treatment, whereas the corresponding pair feed control groups did not show any major changes in hepatic morphology (Fig 1 & Fig 2).

**Figure 1:**
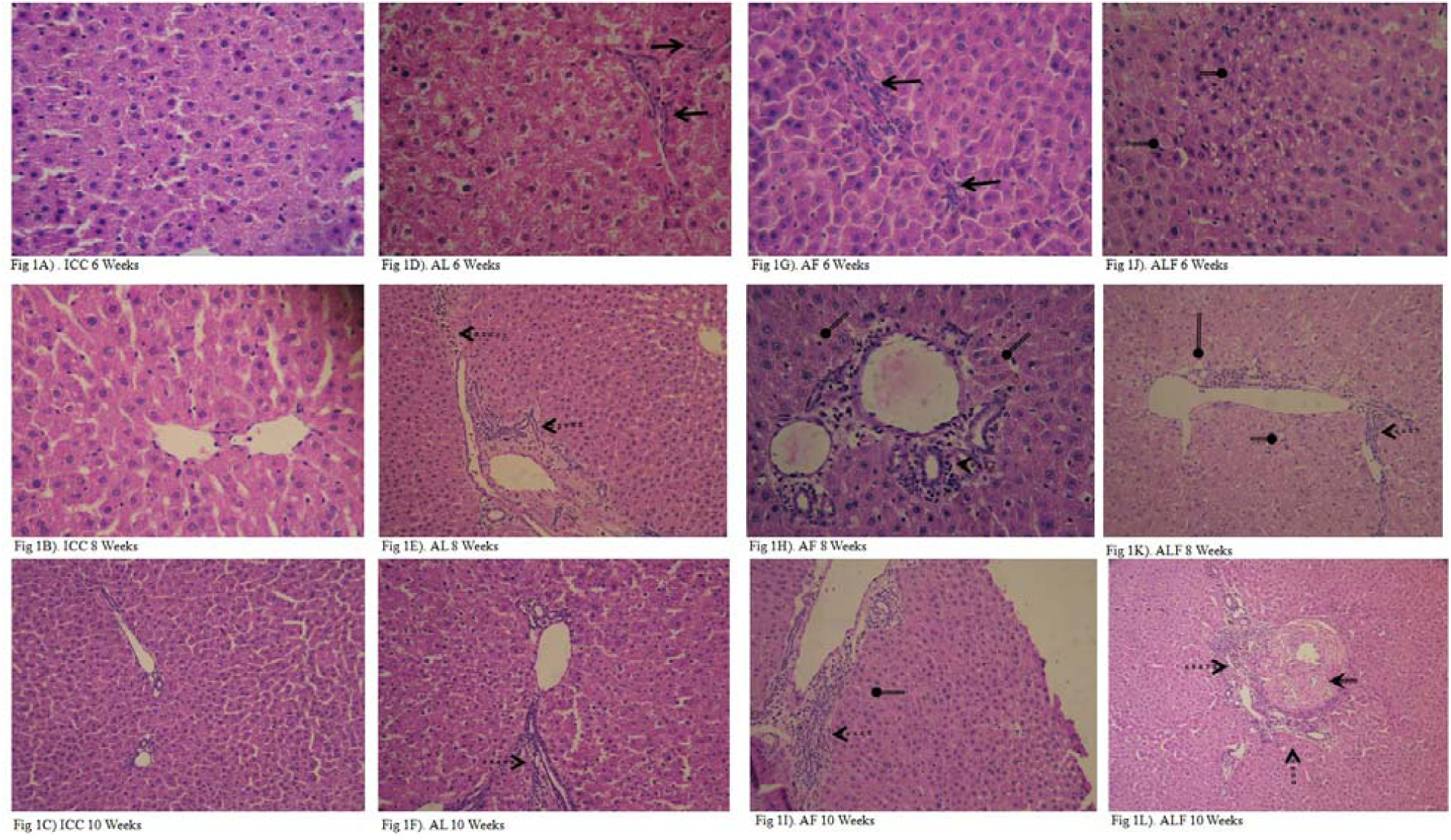
Haemotoxylin & Eosin Staining was performed on 5µm sections of livers taken from different treatment group and these pictures are representative of each treatment group. 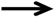 - Focal infiltration of leucocytes in hepatocytes, 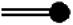 - Leucocyte infiltration in the periportal and perivenous region, 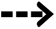 - Hepatic steatosis (microvascicular steatosis, ballooning degeneration), 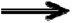 - Granuloma formation.

**Figure 2:**
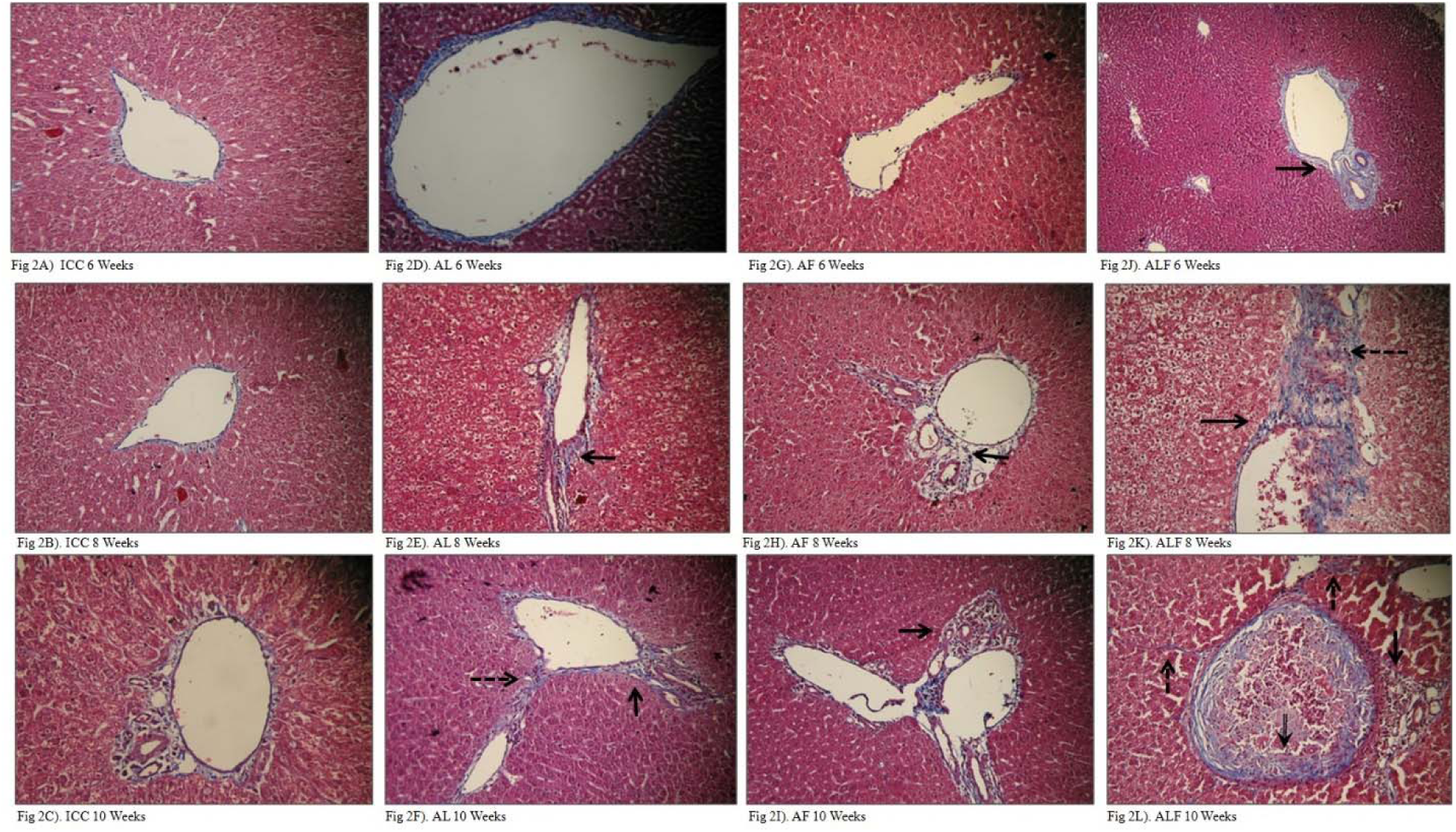
Masson Trichrome Staining was performed on 5µm sections of livers taken from different treatment groups and these pictures are representative of each treatment group. 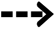 - Pericellular/bridging/chicken wire fibrosis, 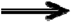 - Fibrosis around the periportal and perivenous region 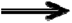 - Loss of liver architect-

H&E staining and Masson trichrome staining were performed to investigate histological changes in liver after chronic alcohol administration. The liver sections from Isocaloric control animals (group 1, 2 & 3) showed normal cords of hepatocytes around the portal track and normal hepatic architecture (Fig 1 A-C). Sinusoidal swelling or infiltration of leukocytes was not observed in these sections. The classical features of hepatopathy (fat droplets or collagen strands) were not evident in liver sections of control group of animals. (Fig 2 A-C, 3A-C)

**Figure 3:**
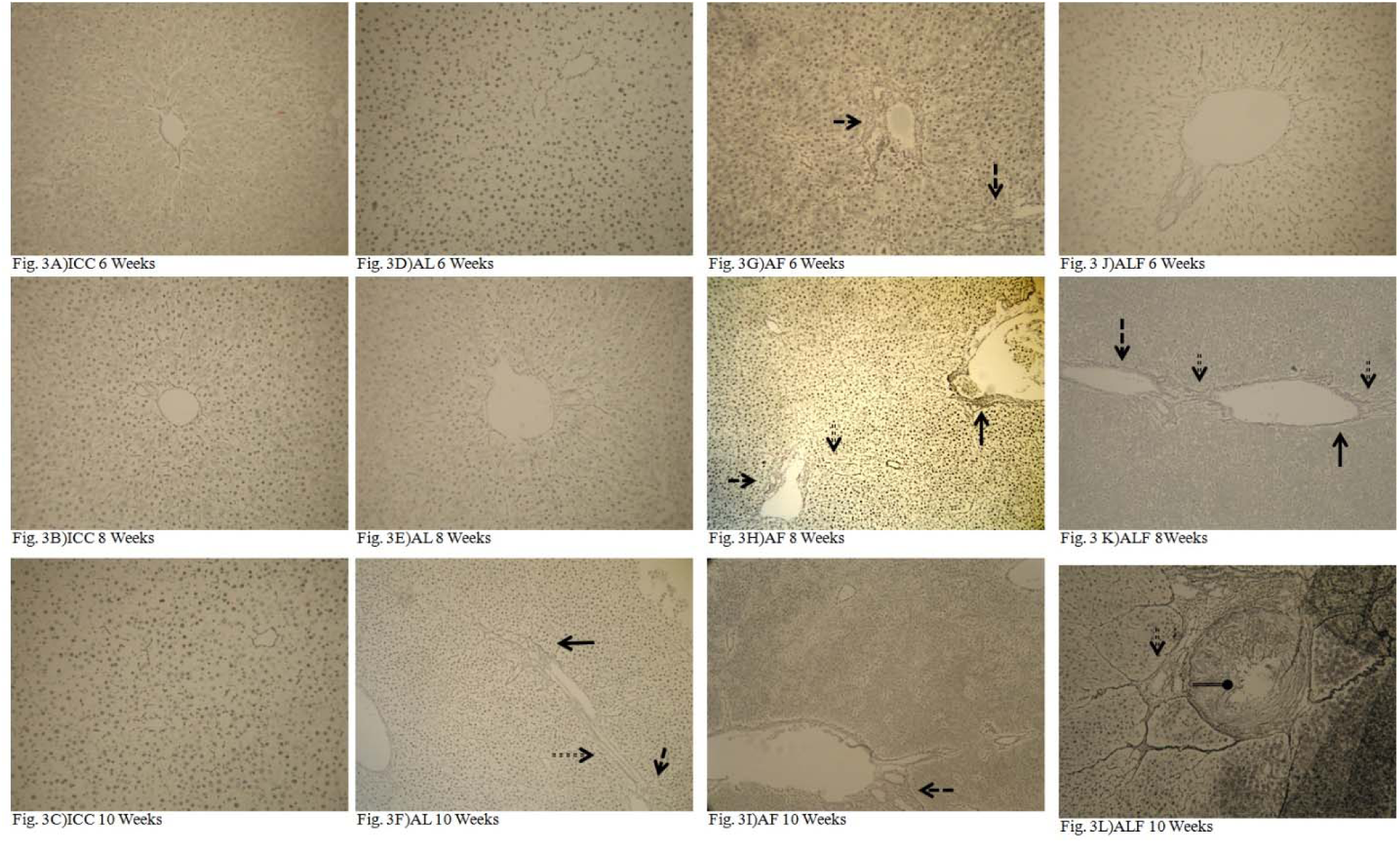
Reticulin Staining was performed on 5µm sections of livers taken from different treatment groups and these pictures are representative of each treatment group. 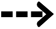 - Pericellular collagen deposition, 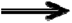 - Collagen deposition around the periportal and perivenous region, 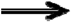 - Loss of liver architect

Liver sections from alcohol and LPS treated animals (group 4, 5 & 6), stained with H&E demonstrated mild degenerative changes following 6 weeks treatment and minimal inflammatory changes were observed after 8-& 10-weeks treatment (Fig. 1 D-F). There were no signs of hepatic fibrosis or steatosis even after 10 weeks of treatment in these groups of animals as observed by masson trichrome and reticulin staining. (Fig 2 D-F, 3D-F)

In contrast when alcohol was administered along with poly unsaturated fatty acids (Corn oil) (group 7, 8 & 9), six weeks treatment showed minimal infiltration in the periportal region with minimal changes in the hepatic architecture in comparison to the corresponding iso caloric control group. Administration of alcohol and fat for eight weeks registered minimal steatosis, moderate inflammation and mild perivenous fibrosis along with minimal collagen deposition. This further progressed to mild steatosis, leukocyte infiltration (Fig 1 G-I) and mild fibrosis.(Groups 7, 8 and 9) (Fig 2 G-I, 3G-I).

The Wistar rats treated with alcohol in conjunction with fat and LPS (Groups 10, 11 and 12) showed progressive increase in hepatic injury. Six weeks treatment resulted in moderate micro-vesicular steatosis and ballooning degeneration along with mild fibrotic changes in the perivenous region and portal tracts. At 8 weeks the hepatic sections displayed moderate leukocyte infiltration and modest fibrosis in perivenous region. After 10 weeks treatment with alcohol, fat and LPS, leukocyte infiltration in portal tracts and perivenous region was observed (Fig 1 J-L). In addition, granuloma formation and loss of hepatic architecture was also evident in the 10 weeks ALF treatment group. Further Masson trichrome staining of these sections demonstrated evident fibrotic changes in periportal, perivenous, pericellular (bridging fibrosis) regions (Fig 2 J-L, 3J-L).The ALF group registered the hallmarks of ALD (steatosis, hepatitis, fibrosis and cirrhosis) with longitudinal progression spanning 6, 8 and 10 weeks.

### AST & ALT Analysis supports graded loss of liver function in ALF treated group

AST and ALT levels are considered as direct measure of liver functions. In the present study various groups of animals were tested for AST and ALT levels in serum samples at stipulated time durations. There was a continuous increase in their levels from six weeks onwards, w.r.t duration of treatment in various groups. In the Isocaloric control groups (Grous 1-3) the ALT levels were 9.5 ± 0.923 U/L; 9.86 ± 0.321 U/L and 10.32 ± 0.325 U/ L after 6, 8 and 10 weeks respectively. The AST levels in the same samples were 11.121 ± 0.33 U/L, 12.131 ± 0.251 U/L; and 13.125 ± 0.4795 U/L respectively. There was no significant difference between the groups (1-3) with respect to treatment duration. In AL and AF treatment groups ALT and AST levels were significantly increased in comparison to isocaloric controls after 8 weeks of treatment. The ALF treatment (group10-12) resulted in significant difference in AST levels evident from the 6th week and the levels increased from 38.875 ± 6.74 U/L (at 6 weeks) to 88.38 ± 13.23 (at 8 weeks), to 172 ± 21.5 U/L (at 10 weeks). The ALT levels increased from 17.5 ± 6.422 U/L (6 weeks), to 28 ± 4.51 (at 8 weeks) to 51.25 ± 8.96 U/L (10 weeks) (Fig 4a-4b).

**Figure 4.**
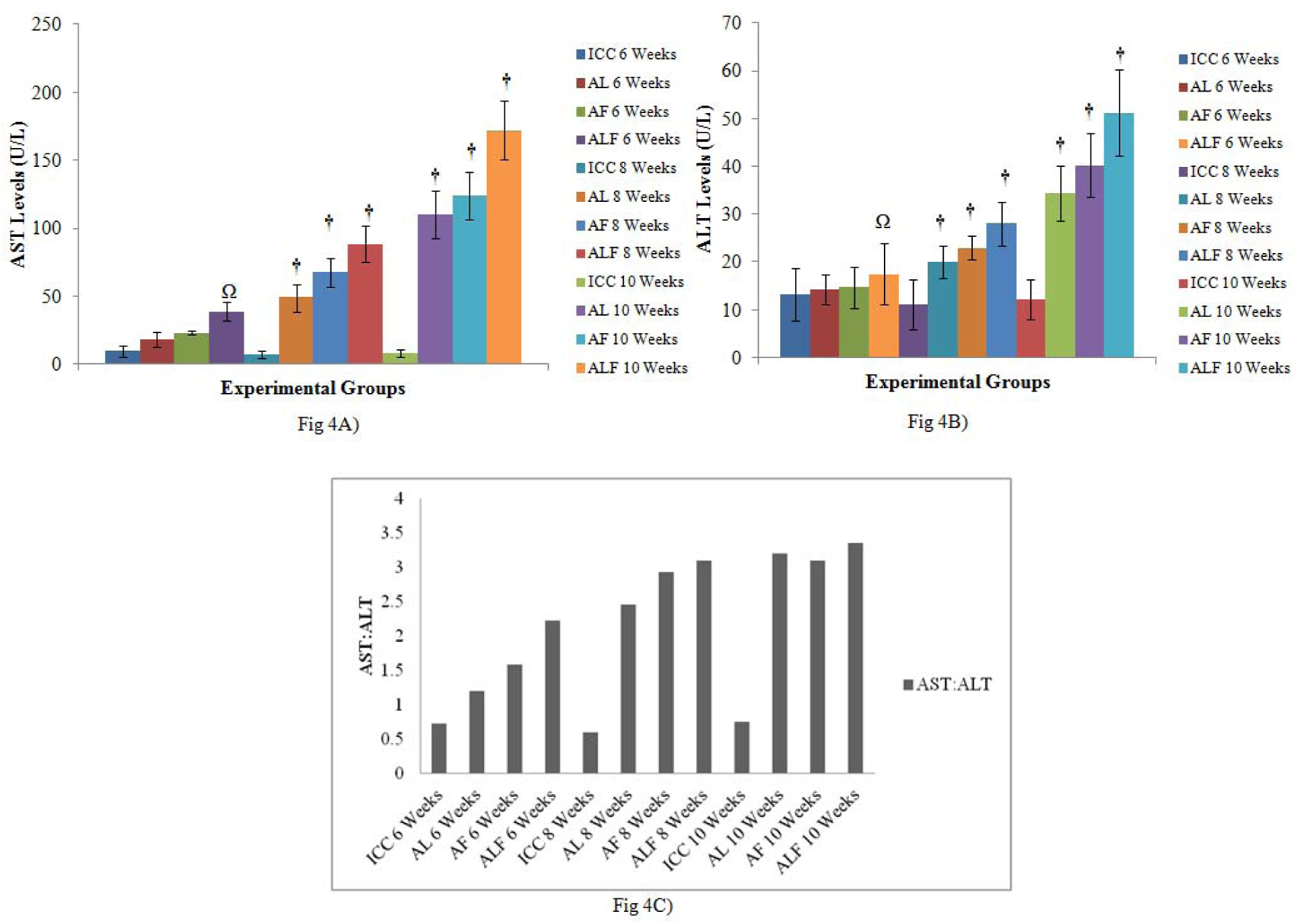
**A) and 4B)** represent the **AST Levels and ALT Levels** observed in serum for all treatment groups. A significant difference between the treated and control group animals appeared earliest in the ALF treatment group. Results are expressed as Mean ± SEM (n=8). † - p<0.001, Ω - p< 0.01, ns - not significant. Figure 4C) shows the ALT to AST ratio for different treatment groups. A typically high ratio falling in range of 2.21:1 to 3.35: 1 in ALF treatment group, represents alcohol related tissue injury.

Moreover, the AST to ALT ratio was more than 2:1 in all the groups particularly after 8 weeks, which is taken as index of disease progression due to alcohol induced damage^14^ (Niemelä et al., 2002) (Fig 4c). In the ALF treatment group the ratio went on increasing from 1.65:1 in 6^th^ weeks to 3.35:1 in 10^th^ week indicating a significant and early derangement of liver functions.

**Blood alcohol content** of the alcohol or alcohol with fat treated wistar rats in the presence or absence of LPS were compared with the isocaloric control groups of animals. Significant and progressive increases in basal and peak alcohol content in plasma were recorded in all treatment groups in comparison to controls (Fig 5a-d).

**Figure 5.**
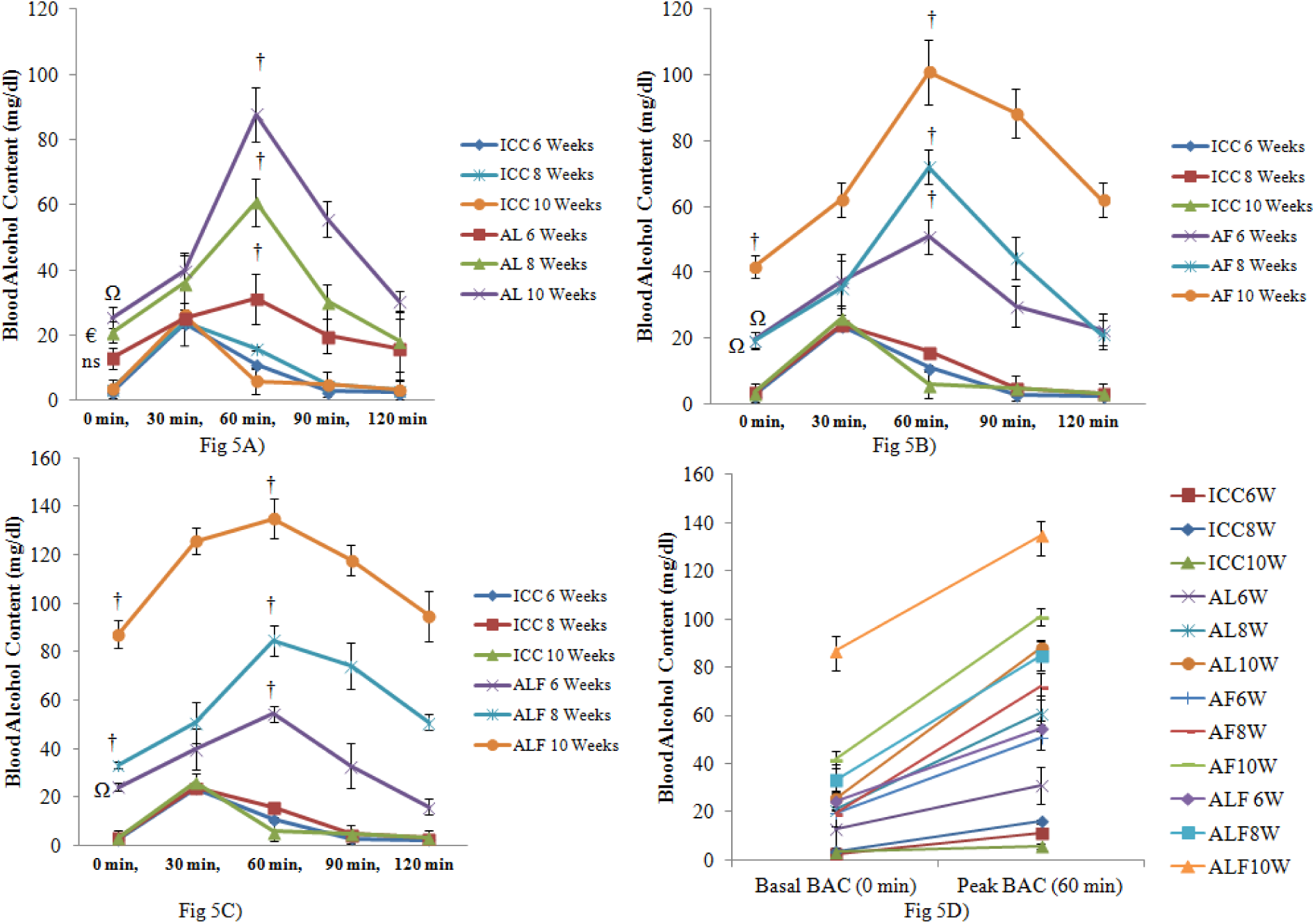
Blood Alcohol Content (mg/dl) of A) AL B) AF & C) ALF group at different treatment durations of 6, 8, & 10 weeks was measured and compared with the ICC group of same treatment duration. Results are expressed as Mean ± SEM (n=8). Figure 5 **D)** Represents the basal and peak levels of BAC in serum of different treatment group animals., €-p<0.05 vs. control, Ω-p<0.01vs. control, -p<0.001vs. control.

In the control group (1-3) the base line alcohol levels (at 0 min) following a single bolus dose of 40% v/v (2g/kg body weight) alcohol were 2.7 ± 0.65 mg/dl, 3.61 ± 0.125 mg/dl & 3.54 ± 2.832 mg/dl whereas peak alcohol levels in plasma were 23.62 ± 0.154 mg/dl, 24.04 ± 0.982 mg/dl & 26.09 ± 3.872 mg/dl (after 60 min) at 6, 8 & 10 weeks respectively. The intergroup differences were not significant (p<0.001).

In the AL treatment group (4-6) **the base line alcohol levels** (at 0 min) following a single bolus dose of 40%v/v (2g/kg body weight) alcohol were 12.99 ± 3.205 mg/dl, 20.98 ± 3.394 mg/dl & 25.33 ± 3.281 mg/dl whereas peak alcohol levels in plasma were 31.5 ± 7.61 mg/dl, 60.9 ± 7.297 mg/dl & 87.79 ± 8.364 mg/dl (after 60 min) at 6, 8 & 10 weeks respectively.

In the AF treatment group (7-9) the base line alcohol levels (at 0 min) following a single bolus dose of 40% v/v (2g/kg body weight) alcohol were 19.48 ± 2.284 mg/dl, 19.32 ± 2.637 mg/dl & 41.8 ± 3.467 mg/dl whereas peak alcohol levels in plasma were 50.91 ± 5.26 mg/dl, 72.07 ± 5.36 mg/dl & 100.98 ± 9.82 mg/dl (after 60 min) at 6, 8 & 10 weeks respectively.

In the ALF treatment group (10-12) the base line alcohol levels (at 0 min) following a single bolus dose of 40% v/v (2g/kg body weight) alcohol were 24.34 ± 1.397 mg/dl, 33.39 ± 1.311 mg/dl & 87.17 ± 5.764 mg/dl whereas peak alcohol levels in plasma were 54.39 ± 3.434 mg/dl, 84.85 ± 6.272 mg/dl & 134.95 ± 8.242 mg/dl (after 60 min) at 6, 8 & 10 weeks respectively. Considerably higher basal BAC in ALF group reflects increased tolerance to alcohol effect.

In the present study we observed a **steady increase in MDA levels** in alcohol treated animals with an increase in treatment duration, which was significantly higher than corresponding control groups (Fig 7). MDA levels in the control group (groups 1-3) at 6, 8 &10 weeks were 22.21 ± 3.351 nmoles/hr/mg of protein; 24.63 ± 4.587 nmoles/hr/mg of protein & 24.86 ± 3.856 nmoles/hr/mg of protein respectively. There was no significant difference within these groups (p<0.001). MDA levels in both AL and AF treatment group (4-6) at 6, 8 &10 weeks increased progressively from However the highest levels of MDA were observed in the ALF treatment group (Fig 6A). This indicates progressive increase in malondialdehyde in liver tissue following chronic alcohol or alcohol and fat administration.

**Figure 6A).**
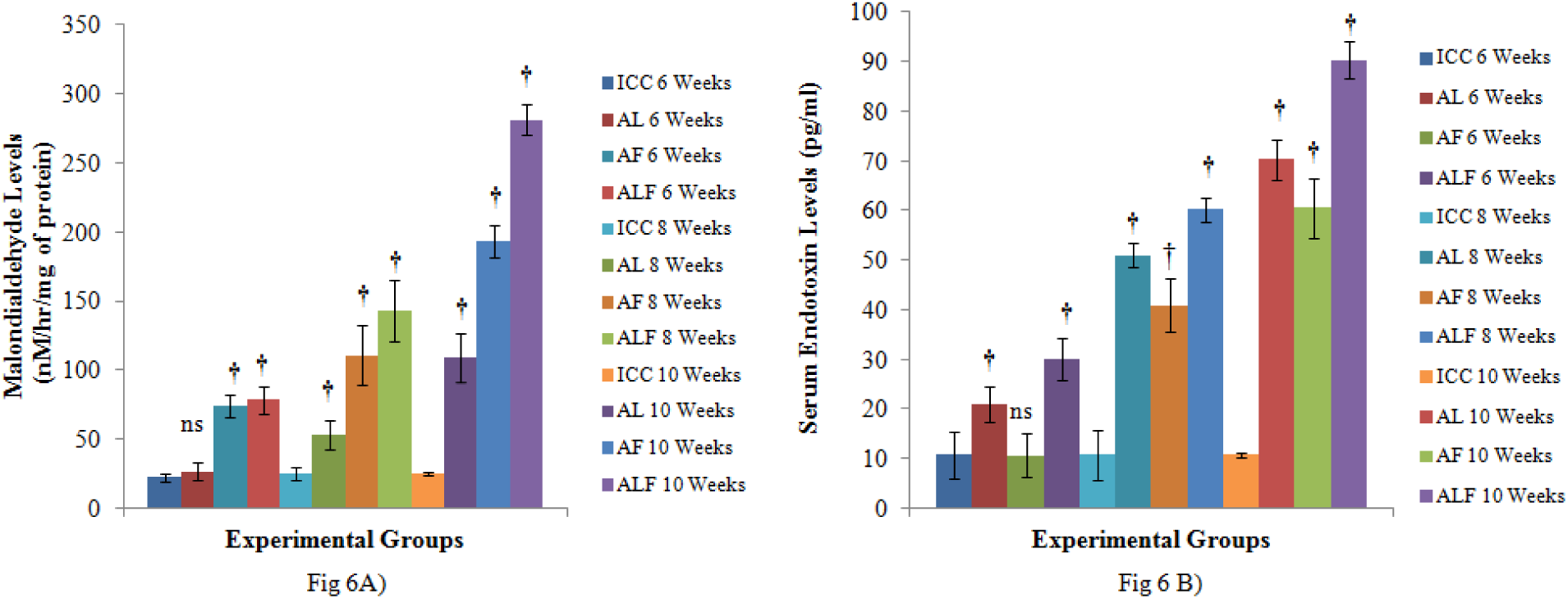
MDA Levels (nm/mg of protein) were taken as measure of lipid peroxidation in tissue. A constant and significant increase in its value represents an increase in lipid peroxidation. Values are represented as Mean ± SEM, (n=8); ns-not significant (p<0.05), - p<0.001vs. control. **Figure B) Serum Endotoxin Levels (pg/ml)** were measured in all treatment groups and results were compared with ICC group of respective treatment duration. Results are expressed as Mean ± SEM (n=8). † - p<0.001, ns – not significant (p<0.05)

**Figure 7.**
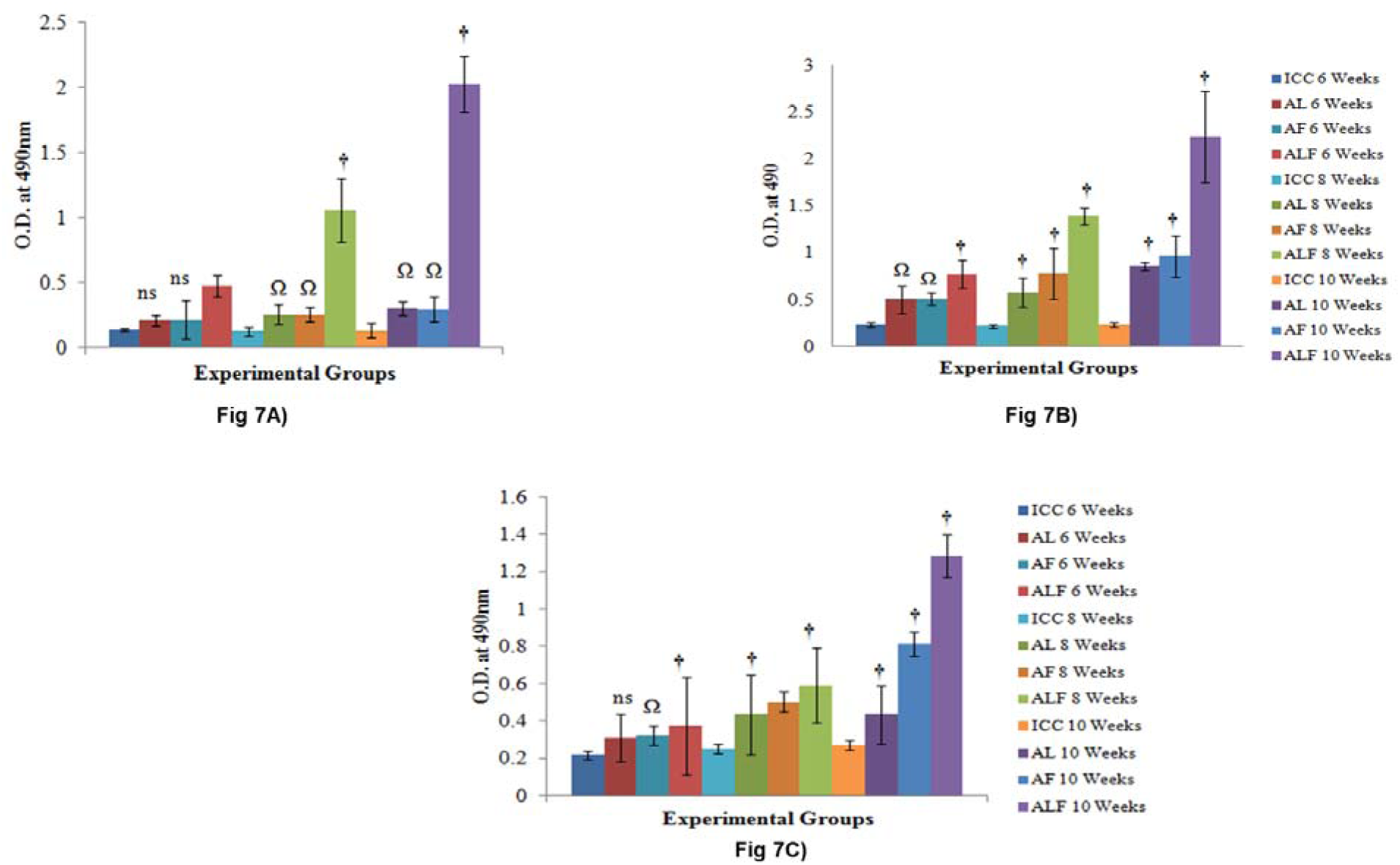
Levels of Antigenic proteins in the Cytosolic fraction of liver proteins ELISA wa performed to estimate the relative levels of **A) IgG B) IgA C) IgE** reactive proteins in all the treatment groups. Values are represented as Mean ± SEM; ns-not significant, €-p<0.05 vs. control, Ω - p<0.01vs. control, - p<0.001vs. control.

**Endotoxin levels** in the serum samples of wistar rats of different treatment groups were compared with the respective isocaloric control groups. We observed a progressive and significant increase in serum endotoxin levels after 6, 8 and 10 weeks of treatment in AL (20.91667± 1.424 pg/ml, 50.91± 1.01 pg/ml, 70.25± 1.688 pg/ml) & ALF (30.12333± 1.7341 pg/ml, 60.133± 1.05 pg/ml, 90.26667± 1.5202 pg/ml) treatment groups w.r.t duration of treatment, whereas in the AF treatment groups the endotoxin levels were comparable to isocaloric controls at six weeks (10.668 ± 1.771 pg/ml) which increased to 40.85± 2.203 pg/ml and 90.266 ± 1.52 pg/ml after eight and ten weeks of alcohol and fat treatment respectively (Fig 6B).

**The antibody responses against cytosolic** liver proteins in the serum was compared using enzyme linked immunosorbent assay (ELISA), the absorbance noted at 490nm was compared between the groups as direct measure of antibody titer. The highest and most significant increase in antibody titer was observed in the ALF treatment group

### IgG Antibody Response

IgG antibody responses against cytosolic liver proteins in the sera of treated and control group of animals was measured. The O.D. at A_490_ in the ICC groups (group1-3) was 0.138±0.029; 0.1243 ± 0.0351; & 0.1299 ±0.0543, at 6; 8; & 10 weeks respectively. In the AL treatment group (group 4-6) the O.D. observed was 0.2078 ± 0.0371; 0.25357 ± 0.0724; & 0.30063 ± 0.05389at 6; 8; & 10 weeks respectively. In the Alcohol + fat treatment group (group 7-9), the O.D. was 0.21333 ± 0.1468; 0.2533 ± 0.0553; & 0.29367 ± 0.0954, at 6; 8; & 10 weeks respectively. The highest O.D. was observed in the ALF treatment groups in a group wide stage specific comparison (<u>group 10-12)</u>. The IgG immune responses against ALF treatment group against cytosolic proteins were 0.47467 ± 0.0836; 1.05133 ± 0.245; & 2.02433 ±0.2134 at 6; 8; & 10 weeks respectively (Fig7A) **IgA Antibody Response** The OD recorded (taken as direct measure of IgA antibody responses) for the cytosolic fractions from liver homogenates of ICC group (group 1-3); were 0.22692± 0.01542, 0.21513± 0.0124, & 0.23407± 0.02318; from AL group (group 4-6) were 0.4997± 0.1504; 0.5762± 0.154; & 0.85267± 0.267; from AF group (group 7-9) were 0.50423± 0.0652; 0.77633± 0.277; & 0.959± 0.2173, and in the ALF group (group 10-12) were 1.38607± 0.0866; & 2.23267±0.483,at 6, 8 & 10 weeks respectively (Fig7B) **IgE Antibody Response** IgE antibody responses against cytosolic liver proteins in the sera of control group animals was 0.21753 ± 0.0236; 0.2531± 0.0262; & 0.2707± 0.238 in AL group was 0.311± 0.127; 0.4337± 0.214; & 0.8108± 0.0521, in the AF group was 0.31957± 0.155; 0.5021± 0.0542; & 0.80187± 0.0633and in the ALF group was 0.37317 ± 0.261; 0.5915 ± 0.20029; & 1.28573± 0.115after 6, 8 & 10 weeks respectively (Fig7C)

### Characterization of Neo-Antigens

In the study we characterized 4 unique proteins in the cytosolic fraction which demonstrated reactivity for IgG (Fig 8A-D), 22 proteins, were identified as IgA reactive (Fig 9A-D), and 12 proteins were identified as immunoreactive for IgE (Fig10 A-D) in the cytosolic fraction of liver proteins harvested from ALF treatment group. Identity of these proteins is given in Table 2,3&4 resp.

**Figure 8.**
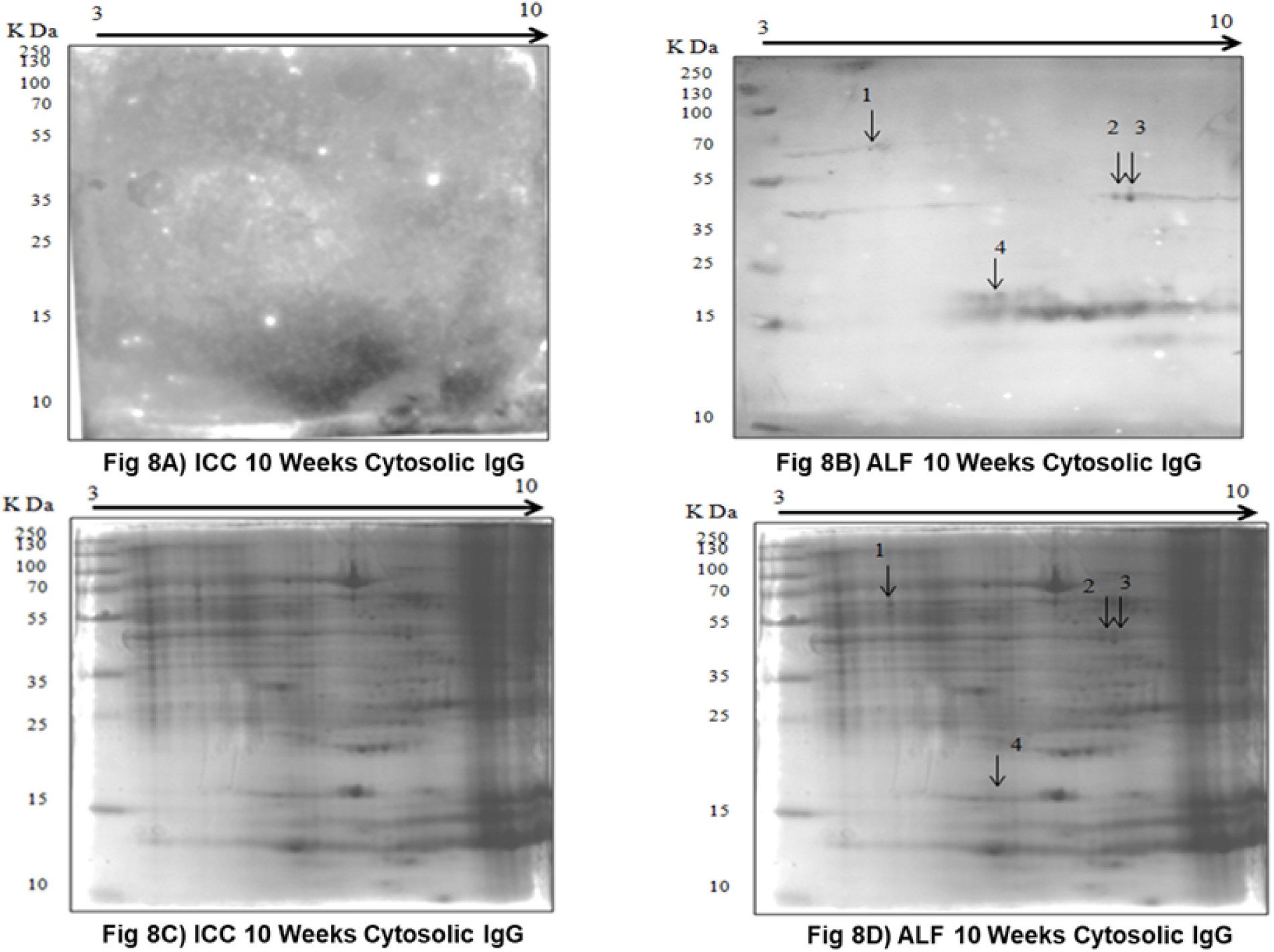
Characterization of IgG reactive proteins in the cytosolic fraction of ALF treatment group. Immunoblot shows no reactivity for IgG in cytosolic fraction of ICC group. Figure 8**-**A, B-Upper panel represents the western blots for cytosolic fraction of ICC and 10 weeks ALF treatment group probed with rat sera of same group (primary antibody) and anti-rat IgG (secondary antibody), Figure 8-C, D-lower panel shows the representative Coomassie stained gels of cytosolic fraction for each treatment duration.

**Figure 9.**
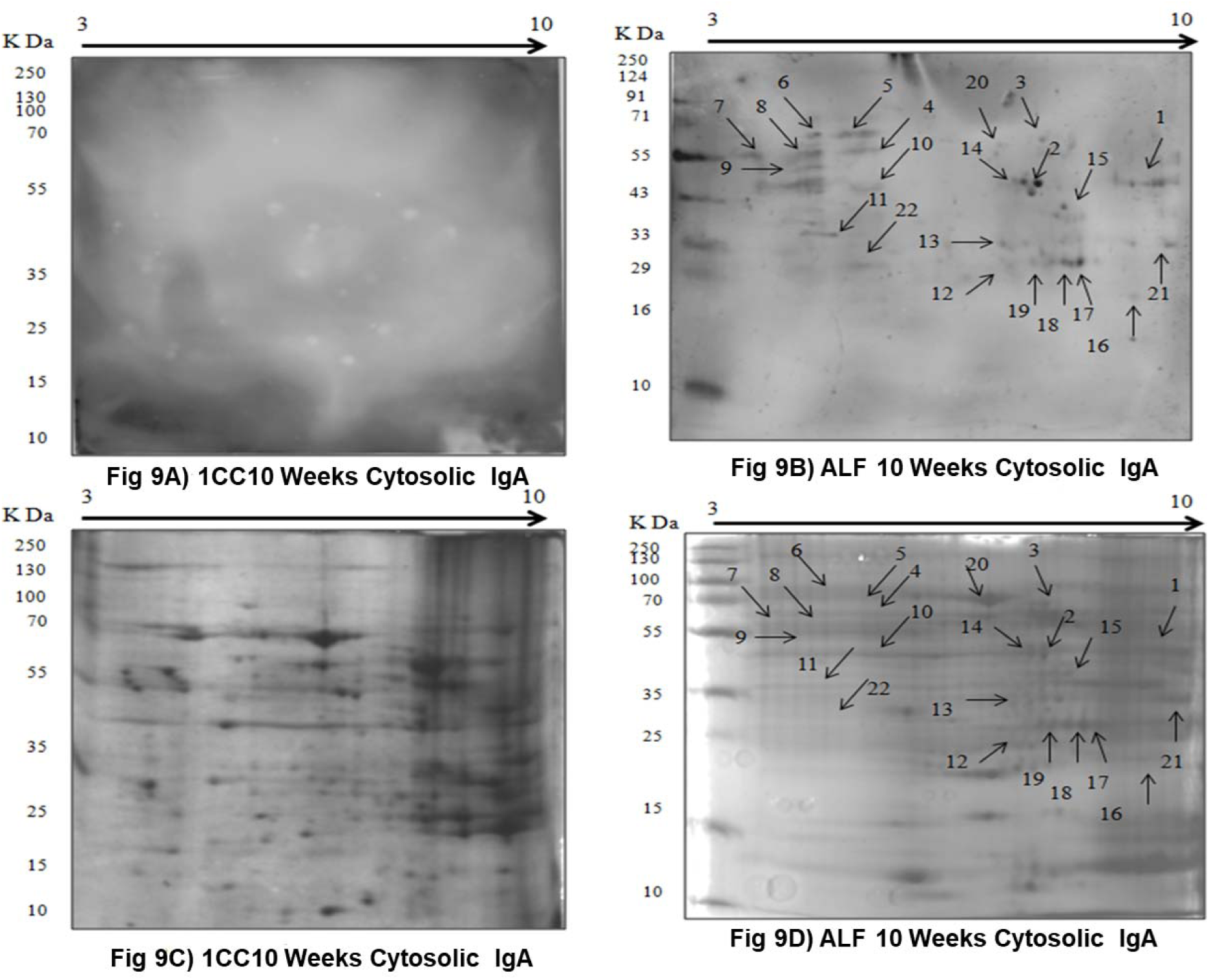
Characterization of IgG reactive proteins in the cytosolic fraction of ALF treatment group. Immunoblot shows no reactivity for IgA in cytosolic fraction of ICC group. Figure 9-A, B-Upper panel represents the western blots for cytosolic fraction of ICC and 10 weeks ALF treatment group probed with rat sera of same group (primary antibody) and anti-rat IgG (secondary antibody), Figure 9-C, D-lower panel shows the representative Coomassie stained gels of cytosolic fraction for each treatment duration.

**Figure 10.**
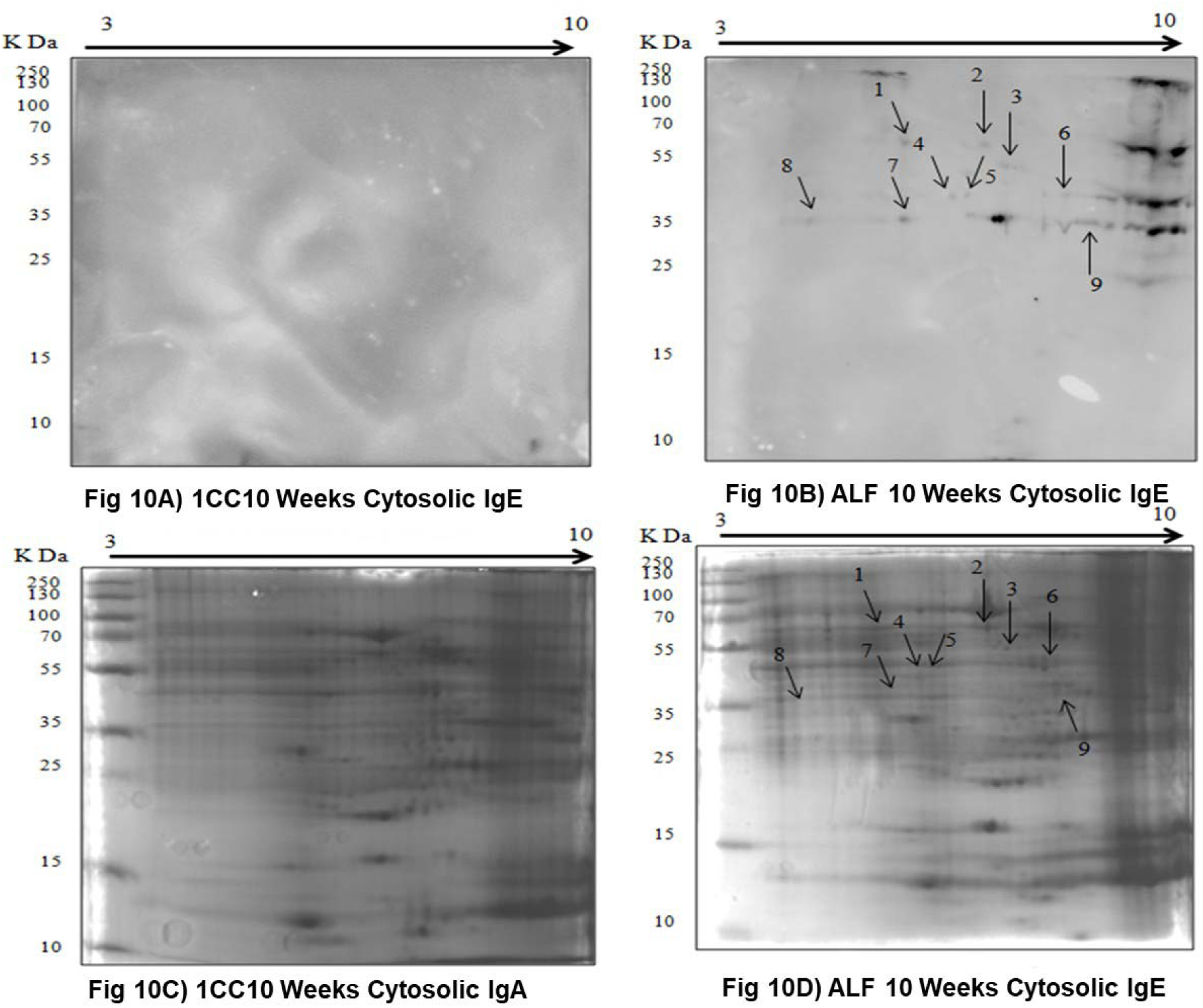
Characterization of IgG reactive proteins in the cytosolic fraction of ALF treatment group. Immunoblot shows IgE reactive proteins in cytosolic fraction of ALF group, Figure 10-A, B-upper panel represents the western blots for cytosolic fraction of ALF treatment group probed with rat sera of same group (primary antibody) and anti-rat IgE (secondary antibody), Figure 10**-**C, D-lower panel shows the representative commassie stained gels of cytosolic fraction for each treatment duration

**Table No. 2:** IgG Reactive Cytosolic Proteins In ALF 10 Weeks Treatment Group.

| Spot No. | Subcellular Location | Accession No. | Protein (Mol. Wt. Kda) | Score | No. of Peptides Matched | Sequence Coverage (%) | No. of Lysine | Biological Function |
| --- | --- | --- | --- | --- | --- | --- | --- | --- |
| 1 | Cytosol | gi25742763 | HSP 70Kda protein 5 Or 78Kda GRP precursor (72.47) | 134 | 19 | 41.74 | 61 | Stress Response, Chaperone |
| 2 | Cytosol<br>(Mitochondria) | gi48675862 | Acetyl CoA thioesterase (49.88) | 82 | 18 | 69.1 | 22 | Fatty Acid Metabolism |
| 3 | Cytosol | gi38649320 | Alpha Enolase (51.74) | 112 | 25 | 65.89 | 36 | Glycolysis, plasmilogen activation |
| 4 | Cytosol | gi818029 | Dismutase (15.74) | 86 | 7 | 43.4 | 9 | Antioxidant |

**Table No. 3:** IgA Reactive Cytosolic Proteins In ALF 10 Weeks Treatment Group.

| Spot No. | Subcellular Location | Accession No. | Protein (Mol. Wt. Kda) | Score | No. of Peptides Matched | Sequence Coverage (%) | No. of Lysine | Biological Function |
| --- | --- | --- | --- | --- | --- | --- | --- | --- |
| 1 | Cytosol<br>(Mitochondria) | gi6981112 | Isovaleryl CoA dehydrogenase (46.86) | 72 | 7 | 25.47 | 23 | Fatty acid $\beta$ oxidation |
| 2 | Cytosol | gi38649320 | Alpha Enolase (51.74) | 112 | 25 | 65.89 | 36 | Glycolysis, plasmilogen activation |
| 3 | Cytosol | gi6978607 | Catalase (60.06) | 157 | 21 | 55.79 | 30 | Redox |
| 4 | Cytosol | gi6980956 | Glutamate dehydrogenase 1 mitochondrial precursor (61.72) | 69 | 19 | 40.32 | 33 | TCA Cycle, Oxidoreductase |
| 5 | Cytosol | gi25742763 | HSP 70Kda protein 5 Or 78Kda GRP precursor (72.47) | 134 | 19 | 41.74 | 61 | Stress Response, Chaperone |
| 6 | Cytosol | gi77917614 | Dynamin 1 like protein (80.41) | 65 | 14 | 38.55 | 43 | Endocytosis, necrosis |
| 7 | Cytosol | gi202549 | iodothyronine 5 monodeiodinase, | 73 | 18 | 47.5 | 51 | Chaperone and Isomerase |
|  | (ER) |  | partial (54.36) |  |  |  |  |  |
| 8 | Cytosol<br>(ER) | gi1352384 | Protein disulfide-isomerase A3 (57.04) | 162 | 25 | 62.57 | 50 | Chaperone and Isomerase |
| 9 | Cytosol | gi11693172 | Calreticulin precursor (48.14) | 74 | 16 | 45.19 | 42 | Calcium binding chaperone |
| 10 | Cytosol | gi8393349 | Fumaryl acetoacetase (46.23) | 92 | 17 | 51.79 | 17 | Amino acid degradation (phenylalanine and tyrosine) |
| 11 | Cytosol | gi20302063 | 3-oxo-5-beta-steroid 4-dehydrogenase (37.64) | 120 | 14 | 48.47 | 25 | Lipid metabolism |
| 12 | Cytosol | gi6978879 | Ferritin heavy chain (21.26) | 78 | 8 | 81.32 | 13 | Iron storage |
| 13 | Cytosol<br>(Mitochondria) | gi57527204 | Electron transport flavoprotein alpha (35.27) | 98 | 15 | 38.14 | 26 | Electron Transport |
| 14 | Cytosol<br>(Mitochondria) | gi48675862 | Acetyl CoA thioesterase (49.88) | 82 | 18 | 69.1 | 22 | Fatty Acid Metabolism |
| 15 | Cytosol | gi6978483 | $\Delta$ aminolavulanic acid dehydratase (36.47) | 91 | 16 | 56.36 | 12 | Heme and Porphyrin biosynthesis |
| 16 | Cytosol | gi8393349 | Glutathione S transferase Yb-3 (25.84) | 60 | 11 | 57.79 | 20 | Glutathione Transfer |
| 17 | Cytosol | gi31377484 | Carbonic anhydrase III (29.43) | 94 | 12 | 62.69 | 18 | Antioxidant |
| 18 | Cytosol | gi31377484 | Carbonic anhydrase III (29.43) | 94 | 12 | 62.69 | 18 | Antioxidant |
| 19 | Cytosol | gi31377484 | Carbonic anhydrase III | 94 | 12 | 62.69 | 18 | Antioxidant |
|  |  |  | (29.43) |  |  |  |  |  |
| 20 | Cytosol | gi12402861<br>2 | Albumin (70.68) | 157 | 15 | 31.41 | 54 | Transport |
| 21 | Cytosol | gi15813854<br>5 | Putative L-aspartate<br>dehydrogenase<br>(30.65) | 80 | 9 | 56.79 | 2 | Pyridine<br>nucleotide<br>biosynthesis |
| 22 | Cytosol | gi15782050<br>9 | potassium<br>channel<br>tetramerisation<br>domain (32.22) | 64 | 11 | 61.48 | 8 | Ion transport |

**Table No. 4:**
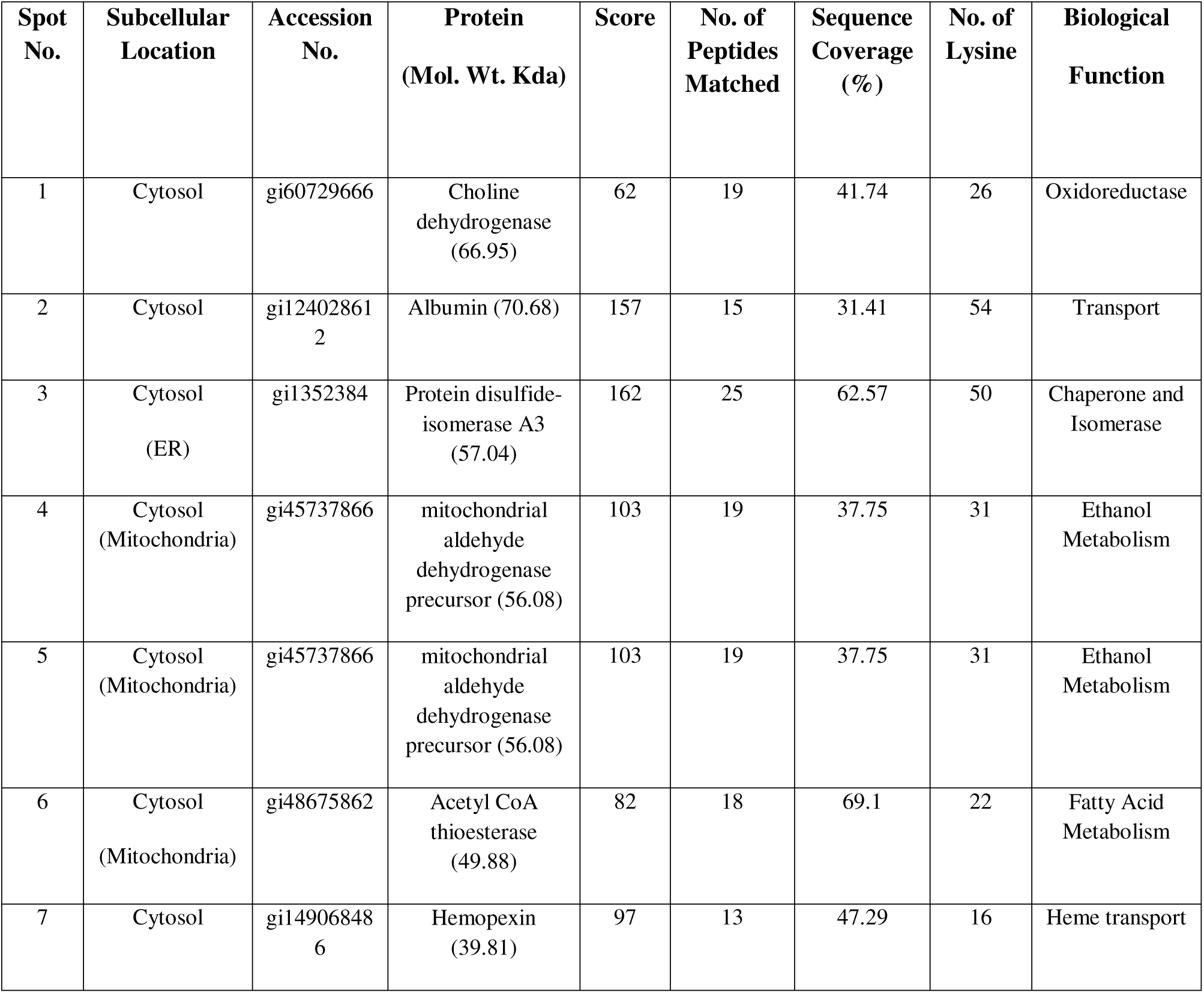
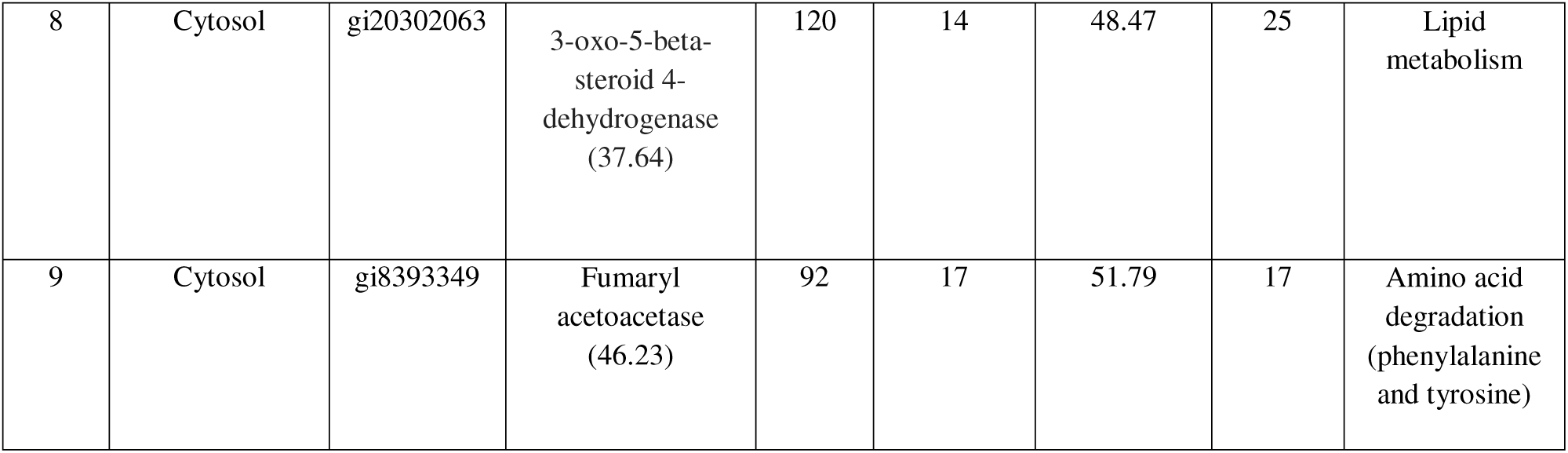
IgE Reactive Cytosolic Proteins In ALF 10 Weeks Treatment Group.

## DISCUSSION

With the evident role of high fat diet and endotoxins in progression of ALD, it becomes essential to employ these two agents in study while addressing the Patho-mechanisms underlying ALD. Therefore, in the current study we co-administered fat (35% of total calories) and LPS (1µg/g body weight) with alcohol (36% to total calories in liquid diet) to develop a comprehensive model of alcoholic liver disease, which progresses through different characteristic stages of the disease. This method is not only easy to perform, but has advantages over previous models, in terms of maintaining consistently high blood alcohol levels, along with marked elevation in AST, ALT levels and of liver histopathology progressing to fibrosis within reasonably short treatment duration of 10 weeks.

The histopathological analysis in our study clearly reveals the potentiating effect of fat & LPS. In the AL treatment group, 10 weeks of treatment lead to only moderate inflammatory changes in perivenous region, without any signs of steatosis or fibrosis, and 6 weeks treatment lead to only mild focal necrosis. In the AF treatment group, the 10 weeks treatment resulted in moderate inflammatory changes and induced moderate hepatic steatosis along-with ballooning degeneration of hepatocytes with moderate fibrosis. However, in the ALF treatment group, mild inflammatory changes were evident within 6 weeks, following 8 weeks of treatment mild steatosis and moderate inflammation along with mild fibrosis were evident in the perivenous region, and 10 weeks of treatment not only lead to moderate inflammation in perivenous and periportal regions but also to granuloma formation along with sever pericellular and periportal fibrosis. Moreover, moderate degree of hepatic steatosis and ballooning degeneration of hepatocytes was also observed. The serum transaminase levels also correlated well with the histological features. We observed a significant increase in AST and ALT levels. The ratio of AST to ALT was observed to be higher than 3:1 in the ALF treatment group following 10 weeks of treatment (3.35:1) which is indicative of alcohol induced liver injury^20, 14^. Further, in the study we were also able to achieve constantly high basal and peak blood alcohol levels in the ALF treatment group. We also observed significant elevation in tissue MDA levels beginning from 6 weeks onwards in all treatment groups, which is indicative of increasing ROS burden in hepatocytes.

In our study we also recorded a significant increase in serum endotoxin levels. Maximum increase in LPS levels was recorded in ALF treatment group treated, in comparison to AL and AF treatment group. Significant increase in serum endotoxin levels was also observed in AF treatment group to which no external endotoxins were administered probably due to alcohol induced enteric dysbiosis and acetaldehyde induced damage to intestinal tight junctions through which endotoxins seep into portal tract. From our results it is suggested that together alcohol, fat and LPS act in a synergistic manner to induce much sever hepatic injury.

Previous experimental reports have shown a direct correlation between high fat diet and pre-disposition to fatty liver ^21, 22^. In the current study also, our results depict a clear potentiation of alcohol induced liver injury by fat and LPS. Furthermore, changes in the biochemical and histological parameters of corresponding ICC, and AF treatment group seen in this study also confirm the symptoms of alcoholic steatosis by the 10^th^ week. These results highlight the role of fat as priming agent. A possible mechanism behind this priming effect could be deposition of excess of lipids in hepatocytes, which not only act as major source of ROS by undergoing peroxidation and generating MDA, but also leads to mitochondrial dysfunction by selectively inhibiting the transport of GSH from cytosol to mitochondria^23, 24^. Aldehydic modification of proteins due to high intracellular MDA levels have also been reported to contribute towards the pathology of disease ^25, 26^. Recent studies also report that excessive lipid deposition in hepatocytes causes disruption of ER homeostasis and upregulation of the apoptotic mechanisms in liver^27,28,29,30^. Another mechanism, by which high fat diet can act as adjunctive insult in development of ALD, is by facilitating direct pouring of adipokines, from the visceral fat, into portal circulation^31^ and simultaneous upregulation of CYP4502E1, which is a prove prominent mechanism for hepatic injury^32^.

The inflammatory responses during ALD have been partly attributed to altered gut barrier permeability due to alcohol metabolites that causes leakage of endotoxins into hepatic portal vein resulting in increased release of circulatory chemokines via CD14/TLR4 pathway^33, 34^. These endotoxins are recognized by LPS binding protein (LBP) and TLR4 receptors^35^ (Kawai et al., 2001) on Kupffer cells (KCs), sinusoidal endothelial cells (SECs) and hepatic stellate cells (HSCs) in liver. Activation of HSCs by LPS leads to increased collagen synthesis, decreased GSH content and upregulation of IL-6^36, 37^. The LPS challenge in ethanol sensitized hepatocytes has been shown to increase secretion of TNF-α, MCP-1 and MIP-2 from SECs, and caspase 3 activity^38, 39, 40^. Simultaneous activation of these pro-inflammatory responses in liver is a potential cause for fibrotic remodeling and necrosis of liver^41^ which is also observed in the current study. In our study, we observed significant increase in leukocyte infiltration as well as in matrix production in the ALF treatment group in comparison to AL and AF treated groups, A partial explanation for this result could be that products of lipid peroxidation (MDA)^42, 43, 44^, and alcohol metabolites sensitize KCs, HSCs, SECs towards LPS by upregulating TLR signaling machinery. This observation supports our initial hypothesis that fat and LPS can act in a synergistic manner to aggravate the innate immune responses, initiating cellular signaling cascade, prompting liver cells towards a necrotic fate^39^.

## Conclusion

Considering the role of dietary fat and endotoxins in disease progression, it is of paramount importance to examine the disease progression in presence of these stimuli. Present study effectively compared the pathological changes using different combinations of ethanol, dietary fat and endotoxin in order to develop an appropriate animal model for alcoholic liver disease. We were successfully able to demonstrate that dietary fat and bacterial endotoxins (lipopolysaccharides) result in potentiation of alcohol induced liver injury. The histopathological changes observed in our experimental model covered the entire spectrum of ALD, as is seen in clinical setting, fatty liver, alcoholic hepatitis, and early stage of cirrhosis, which even the latest accepted model of NIAAA for ALD lacks^45^. We also observed that these animals displayed elevated levels of serum AST, ALT as well as constantly high levels of alcohol were maintained in their blood. Furthermore, the combined treatment of fat, alcohol and LPS lead to striking increase in liver MDA and blood LPS levels, as is observed in clinics. In conclusion, our results indicate that a fairly progressive model for ALD could be developed using this ‘**triple hit’** strategy, where dietary excess of polyunsaturated fat constitutes the first hit, that sensitizes hepatocytes to alcohol and its metabolite which form the directly acting second hit, and finally endotoxin challenge constitutes the third hit, that aggravates the inflammatory responses from various cell types in liver and helps in amplifying the immune responses. Moreover, this new model has 3 critical advantages to its credit (i) easy to set up, (ii) No complicated surgical procedures involved, (iii) effectively replicates the biochemical and histological parameters of ALD as are encountered in clinics till the early cirrhosis stage with reasonably short treatment duration.

## List of Abbreviations

ALD: Alcoholic Liver Disease
ALT: Alanine Aminotransferase
ANOVA: Analysis of Variance
AST: Aspartate Aminotransferase
BAC/BAL: Blood Alcohol Concentration/Blood Alcohol Levels
CPCSEA: Committee for the purpose of Control and Supervision of Experiments on Animals
HSCs: Hepatic Stellate Cells
IFN: Interferon
IKK: IkappaB kinase
IL: Interleukins
IRAK: Interleukin-1 Receptor-Associated kinase
KCs: Kupffer Cells
LBP: LPS Binding Protein
LPS: Lipopolysaccharides
MCP-1: Monocyte Chemotactic Protein-1
MDA: Malondialdehyde
MIP-2: Macrophage Inflammatory Protein 2
MyD88: Myeloid differentiation primary response gene (88) protein
NF-κB: Nuclear Factor kappa B
SECs: Sinusoidal Endothelial Cells
SEM: Standard Error Mean
SGOT: Serum Glutamic Oxaloacetic Transaminase
SGPT: Serum Glutamic-Pyruvic Transaminase
TBK: TANK-binding kinase 1
TLR: Toll Like Receptors
TNF-α: Tumor Necrosis Factor-α
TRIF: Toll/IL-1R domain-containing adaptor inducing IFN.

## Acknowledgement

The authors would like to thank Dr Fouzia Siraj, Pathologist at National Institute of Pathology, Safdarjung Hospital Campus, New Delhi 110029, for her help.

## Author’s Contribution

Shivani Arora, designed, performed and analyzed all the experiments as well as prepared the manuscript; Dr. Anju Katyal critically reviewed the experimental design and the intellectual content of the manuscript; Dr Anju Bansal, an independent pathologist reviewed the slides for pathological features.

## Submission Declaration

The Authors hereby declare that the manuscript submitted for publication has not been previously presented anywhere.

